# MIPSTR: a method for multiplex genotyping of germ-line and somatic STR variation across many individuals

**DOI:** 10.1101/007500

**Authors:** Keisha D. Carlson, Peter H. Sudmant, Maximilian O. Press, Evan E. Eichler, Jay Shendure, Christine Queitsch

## Abstract

Short tandem repeats (STRs) are highly mutable genetic elements that often reside in functional genomic regions. The cumulative evidence of genetic studies on individual STRs suggests that STR variation profoundly affects phenotype and contributes to trait heritability. Despite recent advances in sequencing technology, STR variation has remained largely inaccessible across many individuals compared to single nucleotide variation or copy number variation. STR genotyping with short-read sequence data is confounded by (1) the difficulty of uniquely mapping short, low-complexity reads and (2) the high rate of STR amplification stutter. Here, we present MIPSTR, a robust, scalable, and affordable method that addresses these challenges. MIPSTR uses targeted capture of STR loci by single-molecule Molecular Inversion Probes (smMIPs) and a unique mapping strategy. Targeted capture and mapping strategy resolve the first challenge; the use of single molecule information resolves the second challenge. Unlike previous methods, MIPSTR is capable of distinguishing technical error due to amplification stutter from somatic STR mutations. In proof-of-principle experiments, we use MIPSTR to determine germ-line STR genotypes for 102 STR loci with high accuracy across diverse populations of the plant *A. thaliana.* We show that putatively functional STRs may be identified by deviation from predicted STR variation and by association with quantitative phenotypes. Employing DNA mixing experiments and a mutant deficient in DNA repair, we demonstrate that MIPSTR can detect low-frequency somatic STR variants. MIPSTR is applicable to any organism with a high-quality reference genome and is scalable to genotyping many thousands of STR loci in thousands of individuals.

## Introduction

Variation in short tandem repeats (STRs), which are also known as microsatellites, significantly contributes to phenotypic variation, evolutionary adaptation, and human disease (Gemayel et al. 2012). STRs consist of short (2-10 bp) DNA sequences (units) that are repeated head to tail. The presence of multiple identical or nearly identical adjacent sequence units causes frequent errors in recombination and replication, resulting in loss or gain of units. Consequently, STR mutation rates are 10-10,000 times higher than mutation rates of non-repetitive loci (Eckert and Hile 2009; Legendre et al. 2007).

In spite of their hyper-variability, STRs frequently reside in functional DNA, including coding and regulatory regions. STRs are estimated to be present in six percent of human coding regions (Mularoni et al. 2006; O’Dushlaine et al. 2005), highlighting the potential of STR variation to affect disease risk and other complex traits. Coding STRs that vary among humans tend to reside in genes affecting transcription and neural development (Molla et al. 2009). Several severe genetic diseases, including the trinucleotide expansion disorders Huntington’s and Spinocerebellar Ataxias (SCA), are a consequence of extended STR alleles that act as dominant mutations (Gatchel and Zoghbi 2005). The severity of STR expansion disorders would suggest that natural selection should remove STRs from functional genomic regions, but some, for example the pre-expansion STR allele in SCA2, are maintained by selection (Yu et al. 2005).

Model organism studies have demonstrated significant functional consequences of even subtle unit number variation in select STRs in plants, fungi, flies, voles, dogs, and fish, among other organisms (Fondon and Garner 2004; Hammock and Young 2005; Michael et al. 2007; Rosas et al. 2014; Sawyer et al. 1997; Scarpino et al. 2013; Undurraga et al. 2012). Similarly to humans, STR-containing genes in these organisms tend to be regulatory genes functioning in transcription, development, and sensing environmental factors (Fondon and Garner 2004; Verstrepen et al. 2005). Adding or subtracting a single STR unit can have dramatic phenotypic effects, such as in the polyglutamine-encoding STR in the circadian clock gene *ELF3* in *Arabidopsis thaliana* (Undurraga et al. 2012). STR unit number can show striking non-linear relationships with phenotype, which may in part be due to extensive epistatic interactions with other loci (Butler et al. 2007; Peixoto et al. 1998; Undurraga et al. 2012). Based on existing evidence, STR variation likely comprises an important component of the genotype-phenotype map (*e.g.*, STRs are a viable explanation for some component of the ‘missing heritability’ of genome-wide association studies (Press et al. 2014)), yet due to technological difficulties in genotyping STRs, this component has remained largely undefined.

STRs have almost entirely escaped genome-wide assessment across many individuals due to the complexities of uniquely mapping short, repetitive sequencing reads and the inherently high error rate of STR amplification (*i.e.* amplification stutter). Thus, STR variation is typically excluded or misreported for genomes sequenced with short reads. Recently, several tools have been developed to estimate STR unit number from short read sequencing data (Gymrek et al. 2012; Highnam et al. 2013; Tae et al. 2013). These tools rely on the use of only STR-spanning reads with unique flanking regions to improve mappability and ascertain STR unit number. This restriction imposes size limits (read lengths in extant data are generally 101 bp or less) and greatly reduces coverage of informative reads (**Supplemental Fig. 1**). For example, when assessing the genotype of an STR locus of ∼30 bp for a genome sequenced with 101 bp reads at 5X coverage, one will have to rely on fewer than three STR-spanning reads on average. Moreover, these tools model technical error due to amplification stutter based on STR genotypes from sequenced homozygous or haploid genomes, ignoring the expected diversity of somatic alleles within individuals. These probabilistic models lose applicability in practice, because STR genotype calls are made with as few as one or two STR-spanning reads. Another recent method uses paired-end sequencing reads to infer variation at STR loci, similar to previous methods to detect large insertions and deletions (Chen et al. 2009; Grimm et al. 2013; Hajirasouliha et al. 2010; Qi and Zhao 2011). Due to the resolution limits of gel size selection, this method infers only whether STRs are variable rather than calling STR unit number genotypes (Cao et al. 2014). Thus, the comprehensive assessment of accurate STR genotypes from short-read sequencing data has remained a largely intractable problem.

Vast numbers of genomes, including genomes of hundreds *A. thaliana* strains have been generated with 36 to 64 bp read lengths (Cao et al. 2011; Gan et al. 2011) that are too short for the aforementioned tools. The existing read lengths and coverage depths of these genomes are sufficient to call most single nucleotide variants (SNVs), but insufficient to understand STR variation. It would be inefficient and costly to re-sequence whole genomes of hundreds of individuals or strains with sufficient depth and the longer reads necessary to understand STR variation (∼150-300 bp, >30x coverage) when STRs only make up a small portion of the genome.

The challenges of STR genotyping can be addressed by targeted STR capture to increase the number of STR-spanning reads combined with a sequencing technology that accommodates longer reads to improve mappability and STR genotype calling. Such strategies were recently applied to the human genome, using STR-targeted microarray capture or RNA probe capture prior to sequencing (Duitama et al. 2014; Guilmatre et al. 2013). However, these STR capture methods produced only limited enrichment for STR-containing reads with flanking sequence (2.2% of mappable reads (Guilmatre et al. 2013) and 25% of mappable reads (Duitama et al. 2014) and only marginally improved STR coverage for unit number calls (Table 1).

**Table 1.** Technologies for assessing STR variation by targeted capture and high-throughput sequencing.

| Name | Sequencing and analysis strategy | Accepted coverage<sup>a</sup> | Reported accuracy | Reads mapped to STR targets | Efficiency of mapped reads | STR targets successfully genotyped | Ref. |
| --- | --- | --- | --- | --- | --- | --- | --- |
| Array Capture | Human, Illumina HiSeq, RepeatSeq (Highnam et al. 2013) | 2 reads | 88%-92% | 38.7% | 6.5% informative reads | >= 1 genotype for 54.5% of targets across 8 individuals | (Guilmatre et al. 2013) |
| SureSelect RNA probe capture | Human, Roche 454 , locally align flanking regions | 4 reads | 88%-95% | ~60% | 40% informative reads | 30.1%-36.8% of targets per sample | (Duitama et al. 2014) |
| MIPSTR-smMIP capture | *A. thaliana*, Illumina MiSeq, map to locus-specific synthetic reference | 4 reads | 94%-98% | 72% | 55%-64% informative reads | 64% of targets across samples, at least 50% of targets in 90% of samples |  |

a: Minimum coverage of a single STR that is considered sufficient to call a genotype.

Here, we address the major obstacles of STR genotyping with a robust, scalable, and inexpensive method, MIPSTR. MIPSTR combines STR capture via single-molecule molecular inversion probes (smMIPs) (Hiatt et al. 2013) with mid-size sequencing reads and a unique mapping strategy. In proof-of-principle experiments, we captured and sequenced STRs genome-wide in diverse *A. thaliana* populations, called germ-line STR genotypes with high accuracy, and quantified technical error with single-molecule information. Moreover, enabled by single-molecule degenerate sequence tags, we demonstrate that MIPSTR can capture the same STR locus from thousands of different cells, thereby enabling detection of somatic STR variants with high sensitivity.

## Results

### Single molecule capture strategy yields highly accurate STR germ-line genotypes

We employed single molecule Molecular Inversion Probes (smMIPs) (Hiatt et al. 2013) to capture STRs, thereby maximizing the number of STR-spanning, informative reads. In a proof-of-principle experiment, we targeted 102 STRs across the genome of the model plant *A. thaliana*, including exonic, intronic, regulatory (AM Sullivan, AA Arsovski, J Lempe, KL Bubb, MT Weirauch, PJ Sabo, R Sandstrom, RE Thurman, S Neph, AP Reynolds, et al., in press), and intergenic tri-and hexa-nucleotide STRs (**Supplemental Fig. 2, Supplemental Table 1**). We first applied MIPSTR to the reference *A. thaliana* strain Columbia-0 (Col-0), which has been Sanger-sequenced and for which accurate STR genotypes are available for comparison.

For each targeted STR, we designed a MIP, which is an 80bp oligonucleotide that contains: i) targeting arms which will uniquely hybridize to STR flanking regions, ii) a 12bp degenerate tag to distinguish individual capture events, and iii) a common backbone for PCR and sequencing priming (Fig. 1A) (Hiatt et al. 2013). In Col-0, we successfully captured all 102 STR target loci (**Supplemental Fig. 3**). After capture, MIPs were amplified for subsequent sequencing. As STR amplification is prone to PCR stutter and rampant technical error, we performed optimizations including modifying amplification conditions, specifically adjusting extension time, extension temperature, and polymerases used (see Methods).

**Figure 1.**
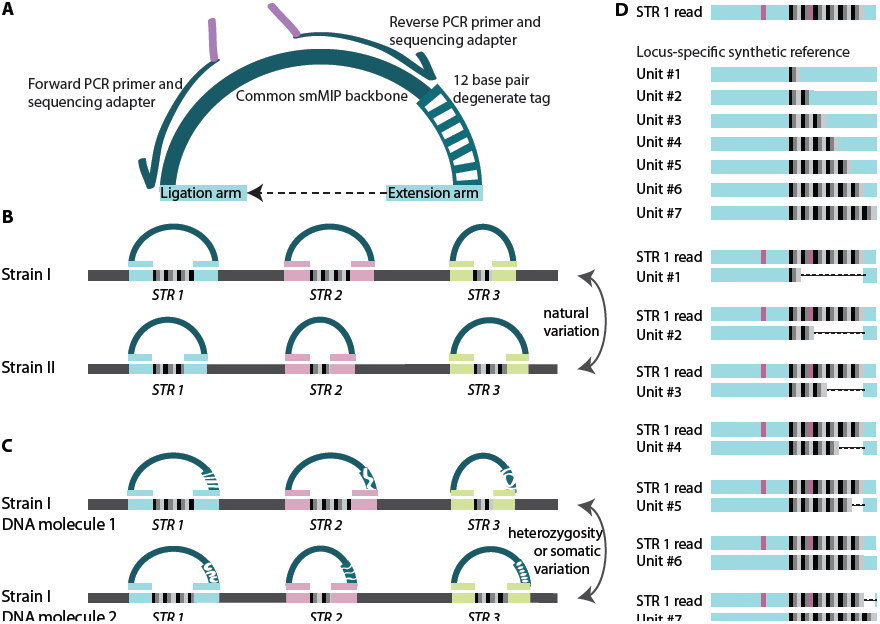
MIPSTR determines germ-line and somatic STR variation with a combination of targeted capture, sequencing, and a novel mapping strategy. **A)** single-molecule molecular inversion probe (smMIP) with common backbone for PCR primer binding (dark-green, also shown PCR and sequencing primers with arrows and purple sequencing adapter), 12 base pair degenerate tag (striped, green/white), and targeting arms with locus-specific, STR-flanking sequence (blue). As shown, one targeting arm is the primer for polymerase extension (extension arm), ligation closes the circle at the other targeting arm (ligation arm). **B**) Applying across individuals identifies germ-line STR variation across genetically diverse individuals. **C**) Applying MIPSTR distinguishes somatic STR variation from technical error, using many degenerate tags (see in **A**). STR variation within a tag-defined read group (*i.e.* reads with the same degenerate tag) is considered technical error. STR variation across tag-defined read groups is considered somatic variation. **D**) MIPSTR maps reads from a given STR locus (based on targeting arm sequence) to locus-specific synthetic references with unit number 1 through 100 (1 through 7 shown here). SNVs (in pink), even if occurring in the STR sequence, do not affect mapping or STR unit number genotype calls.

MIPSTR libraries were sequenced using 250bp forward reads paired with 50bp reverse reads on the Illumina MiSeq platform. The 250bp forward reads spanned the ∼20 bp ligation targeting arm followed by 200bp of target sequence (STR sequence and unique flanking sequence) and ∼20 bp extension targeting arm (large STR expansions will be missing some or all of the extension targeting arm). MIPSTR can assess STRs up to ∼180 bp in length, considerably longer than the STRs currently assessed from whole-genome-sequencing data. The 50 bp reverse reads spanned the 12 bp degenerate tag, which identifies each specific MIP molecule, and the extension targeting arm (Fig. 1A). This experimental design allows MIPSTR to omit the computationally costly and error-prone step of mapping repetitive reads of low complexity to whole genomes. We sorted reads according to their MIP targeting arms, and for each MIP, used BWA (Li and Durbin 2009) to map its corresponding reads to a set of synthetic reference sequences designed specifically for each targeted STR (Fig. 1D). These synthetic references consisted of the STR sequence from the Col-0 reference genome with all possible STR unit number alleles between 1 and 100, which suffices for STR alleles within our size range. We successfully mapped 72% of all sequencing reads to the targeted loci (Table 1).

We called a genotype for each mapped read according to the quality of its alignment to an STR allele sequence (BWA alignment scores >= 180 were called as genotypes). Due to our mapping strategy, variation outside of the STR or SNVs within the STR does not affect STR unit number genotype calls (Fig. 1D). For Col-0, 55% of our mappable reads yielded informative STR unit number calls. Relative to previously described methods, this result represents a dramatic improvement in the number of informative reads per unit of sequencing effort (Table 1), such that it represents a substantial improvement in the efficiency and accuracy of STR genotyping. We required at least four STR-spanning reads at each locus to call an STR genotype. Ultimately, we called unit number genotypes for 96 out of the 102 examined STR target loci. For these loci, our calls were 96% concordant with the Col-0 reference allele, including the highly variable coding STR in the gene *ELF3* (Fig. 2) (Undurraga et al. 2012).

**Figure 2.**
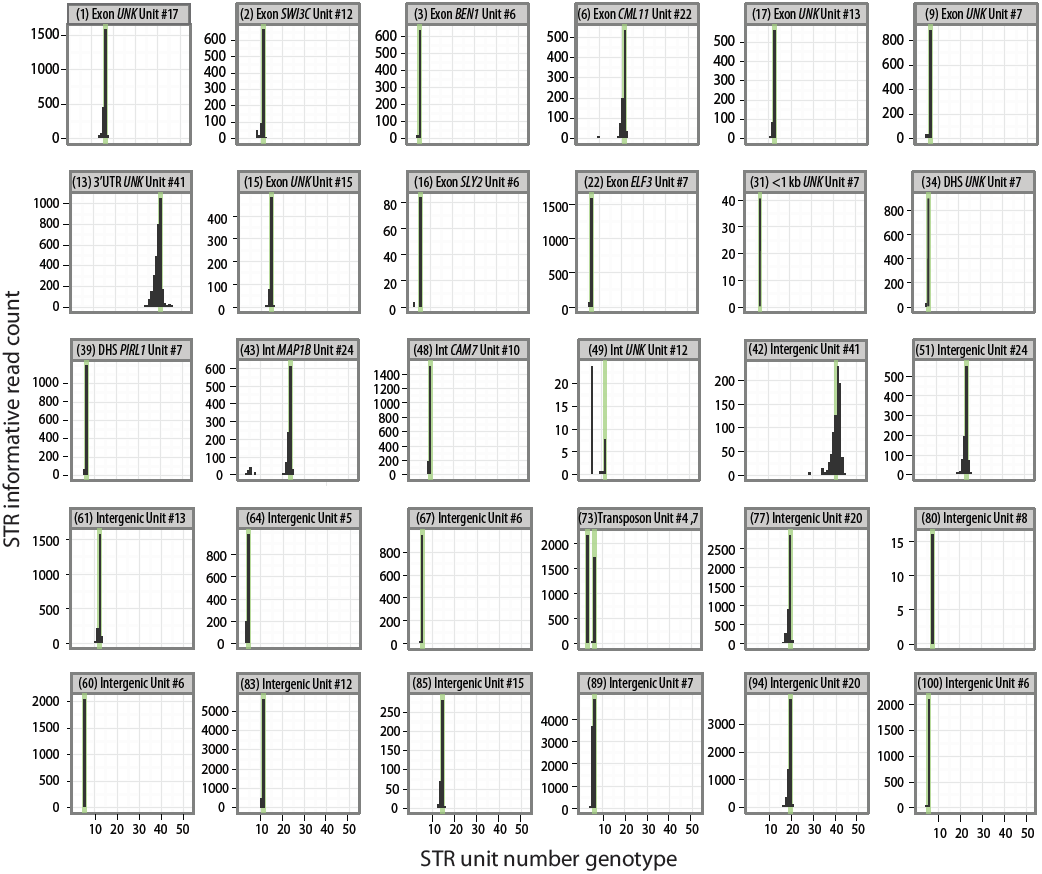
MIPSTR accurately determined germ-line STR unit number in the reference strain Col-0. Raw read counts at 30 representative STR loci, with reference genome STR unit number indicated in green. *UNK* indicates gene of unknown function. Numbers shown in parentheses refer to STR IDs (see Supplemental Table 1). Two instances of genomic duplication (residing in transposons) are shown (STR ID 73 and 89) – both alleles showed comparable read count. Note that erroneous calls show low read count or high technical error.

Most importantly, unlike any previous method that we are aware of, each STR is represented by many independent capture events of STR loci at the pre-amplification stage. Although amplification introduces technical error, MIPSTR distinguishes between technical error, heterozygosity, and somatic mutations by comparing reads within and between capture events (Fig. 1C). The assessment of independent capture events is enabled by the use of smMIPs with degenerate tags (Hiatt et al. 2013), *i.e.* the same STR locus from many different cells is captured by many differently tagged MIP molecules. For each tag-defined read group (*i.e.* reads containing the same degenerate MIP tag), we assumed that the mode of called unit numbers across reads is the true allele for this capture event (Fig. 3). STR unit number variation within a tag-defined read group is considered technical error (Fig. 3, Supplemental Table 1). However, unit number variation observed among different MIP molecules, each representing independent capture events, is potentially the result of heterozygosity, somatic variation, or duplication (Figs. 1C, 3). Using the additional information of tag-defined read groups resolves the distribution of total read counts (compare Fig. 3A to 3B, C) and greatly improves confidence in STR genotype calls. Using information from tag-defined read groups also identified STR loci with consistently high technical error (Fig. 3, middle panel, **Supplemental Table 1**), which can be excluded in subsequent analyses. Furthermore, using information from tag-defined read groups has the potential to detect multiple STR alleles within a single individual (Fig. 3, right panel).

**Figure 3.**
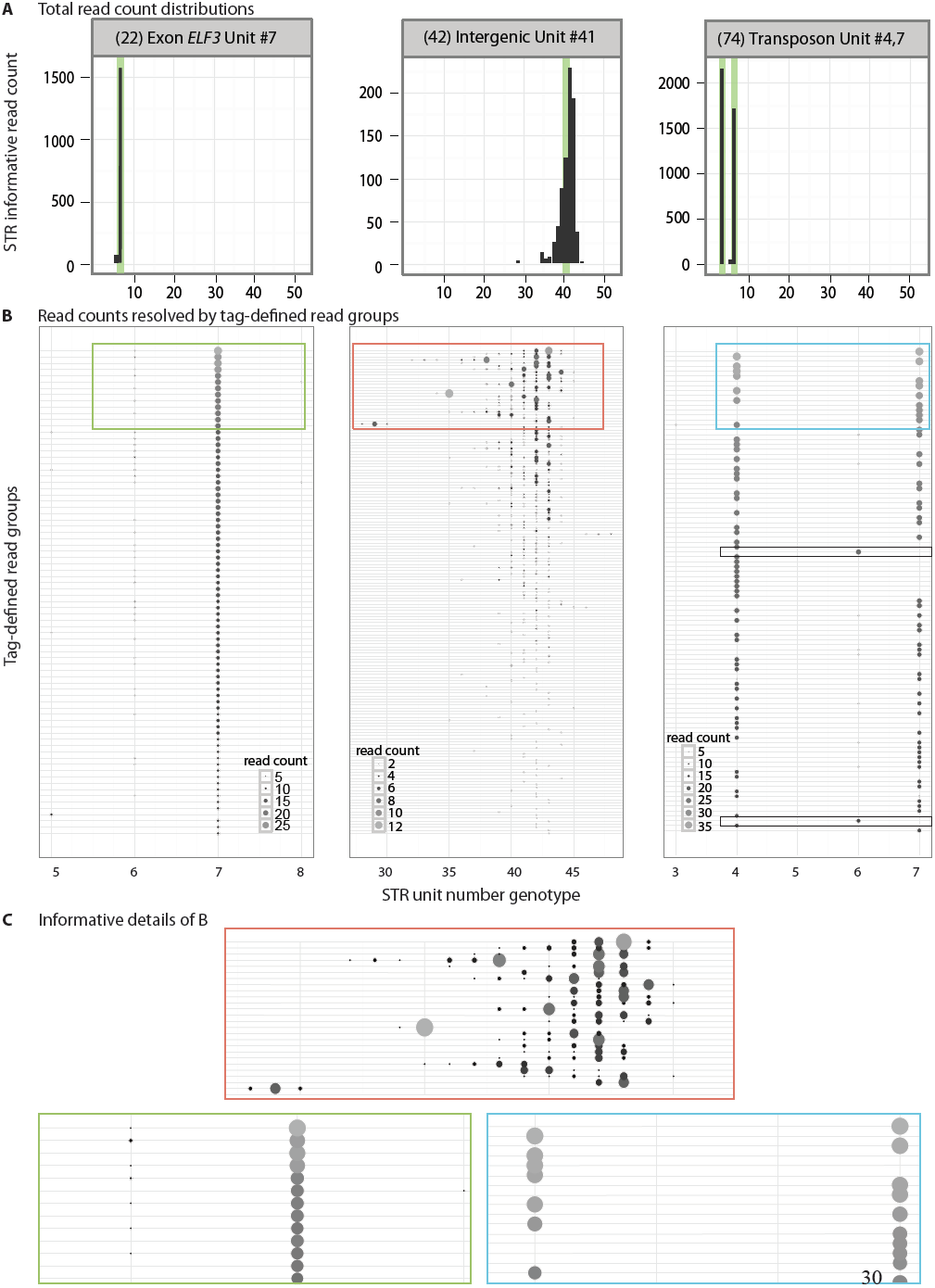
MIPSTR distinguished technical error from somatic variation. **A)** Three histograms from Figure 2 with total read counts. Left, the known *ELF3*-STR unit number is clearly supported by the modal unit number. Middle, this intergenic STR showed great variation in STR unit number; the mode did not support the known STR unit number. Right, this STR resides in two copies in two different genomic locations (transposons). Both known alleles were identified, yet total read counts alone cannot distinguish genomic duplicates from technical or somatic error. **B**) Reads are separated into tag-defined read groups with dot sizes and color representing read count (different scales for each locus, see inset). Colored boxes are shown in detail in C. Left, all tag-defined read groups with one exception supported the known STR unit number seven. Most tag-defined read groups showed low levels of technical error, primarily reads with unit number six (-1), but also five and eight. Middle, separating reads into tag-defined read groups illustrates the extremely high technical error for this STR. The mode of a tag-defined read group was often supported by less than 50% of total reads. Some tag-defined read groups contained as many as six different STR genotypes. We exclude such loci from the analysis of somatic STR variation. Right, as expected for a duplicate STR or a heterozygote, approximately half of the tag-defined read groups support each of the known STR genotypes with very little technical error. We also observed evidence of a somatic STR allele with unit number six, which was supported by two tag-defined read groups (boxed, black outline). Note the absence of either known STR allele for these tag-defined read groups. This STR genotype is also visible in the total read count histogram (**A**, right), where it would be interpreted as technical error by other methods. **C**) Detailed views of plots in B; outline color corresponds to respective plot.

*A. thaliana* is an inbreeding plant and hence assumed to be homozygous at the vast majority of loci. Therefore, to test the potential of our method to detect multiple high-frequency alleles of the same STR, we assessed two STR loci present in two nearly identical copies on two different chromosomes in the Col-0 reference genome. For both STRs, the two genomic copies have different STR unit number genotypes in addition to SNV variation, enabling us to readily distinguish them. Indeed, for both STRs, we detected both unit numbers at high levels.

Specifically, for the STR (STR ID 73a and b) with only one SNV difference between duplicate copies, we observed near equal representation of both alleles (Fig. 3, right panel). We also observed two tag-defined read groups supporting unit number six, which may represent a somatic STR variant in this individual. Without differentiating tag-defined read groups, reads representing this STR genotype would be interpreted as technical error, like the few reads representing ELF3 STR unit number as six (compare Fig. 3 left panel to right panel). This example demonstrates the importance of including single-molecule information in STR genotype analysis.

Furthermore, we found evidence for the duplication of an intergenic STR that is located amidst multiple transposons; this duplication is not present in the Col-0 reference assembly (Fig. 2, STR ID 89). As for the other duplicated STRs, the two alleles, in this case six and seven, were supported by approximately equal number of tag-defined read groups in multiple Col-0 siblings. These results suggest that MIPSTR can readily identify heterozygous and somatic STR variants, which have been largely inaccessible by previous analytical or empirical methods (Gymrek et al. 2012; Willems et al. 2014; Guilmatre et al. 2013; Highnam et al. 2013; Duitama et al. 2014).

## MIPSTR accurately determines STR unit number genotypes across diverse A. thaliana strains

We applied MIPSTR to 96 genetically diverse strains of *A. thaliana*. These strains have been assessed for over 100 quantitative phenotypes and have been previously sequenced, primarily with 36 to 64 bp reads at a coverage of ∼20X, to detect SNVs and structural variation (Cao et al. 2011; Gan et al. 2011). STRs evolve on a different time scale than SNVs, so linkage disequilibrium between STRs and SNVs breaks down quickly (Willems et al. 2014). Therefore, we cannot use linked SNV data to understand the relationship between STR unit number genotype and phenotype. Given the strong potential of STR variation to cause phenotypic variation, we set out to call STR genotypes across many divergent individuals and to show how even data for only 100 STR loci can improve our understanding of the genotype-phenotype map.

We determined the genotypes of the 100 STRs across the 96 diverse strains of *A. thaliana* including the reference strain Col-0 for a total of 9600 targeted STR loci in one Illumina MiSeq v2 sequencing run. MIPSTR scaled well to this task; both the number of targeted loci and the number of examined genomes can be readily increased by several orders of magnitude. STRs tend to be surrounded by repetitive sequence and AT rich regions, but in spite of this challenge, we successfully captured STR loci genome-wide for these genetically divergent strains. Specifically, we captured at least 50 STR loci in 86 out of 96 strains (90%, Table 1) and at least 75 STR loci in 59/96 strains (61%).

To apply MIPSTR to multiple strains, we pooled the 96 strain-specific capture libraries, each with a unique strain barcode on the reverse PCR primer, and sequenced as described above. For these pooled libraries, we sorted reads first by strain-specific barcode, then by targeting arm to identify the STR locus and degenerate MIP tag to identify reads originating from the same capture event (Fig. 1B). Similarly to our results with the reference strain Col-0, we were able to map 72% of sequence reads to their STR target loci and of those 64% were informative for calling STR unit number genotypes (Table 1). In this experiment, the Col-0 library represented ∼1% of the total sequence reads, which should greatly reduce the information for each STR compared to our single Col-0 library run. Despite this dramatic reduction in information content, we could accurately call germ-line STR unit number genotypes for 97% of loci (64 out of 66 loci with at least 4 STR-spanning reads) (**Supplemental Table 2**). Comparing MIPSTR calls for the *ELF3-*STR to genotype calls from previous Sanger sequencing (Undurraga et al. 2012), MIPSTR performed with 98% accuracy (51 out of 52 strains) (Fig. 4). As previously discussed, using information from tag-defined read groups aided us in resolving STR genotypes. For example, for the strain Kin-0, total counts supported unit number 18 and 19 for the *ELF3* STR (Fig. 4). Resolving read counts by tag-defined read groups enabled us to eliminate technical error and call 19 units as the correct Kin-0 *ELF3* STR unit number. Across all 96 strains, we called STR unit number for 60% or more of STR loci in 62% of strains, with a total of 6,179 STR unit number genotypes (out of 9,600 targets or about 64% of targets) determined with a single Illumina MiSeq v2 sequencing run. As previously shown, additional sequencing is expected to yield many more capture events and thus more complete coverage across STRs (Turner et al. 2009).

**Figure 4.**
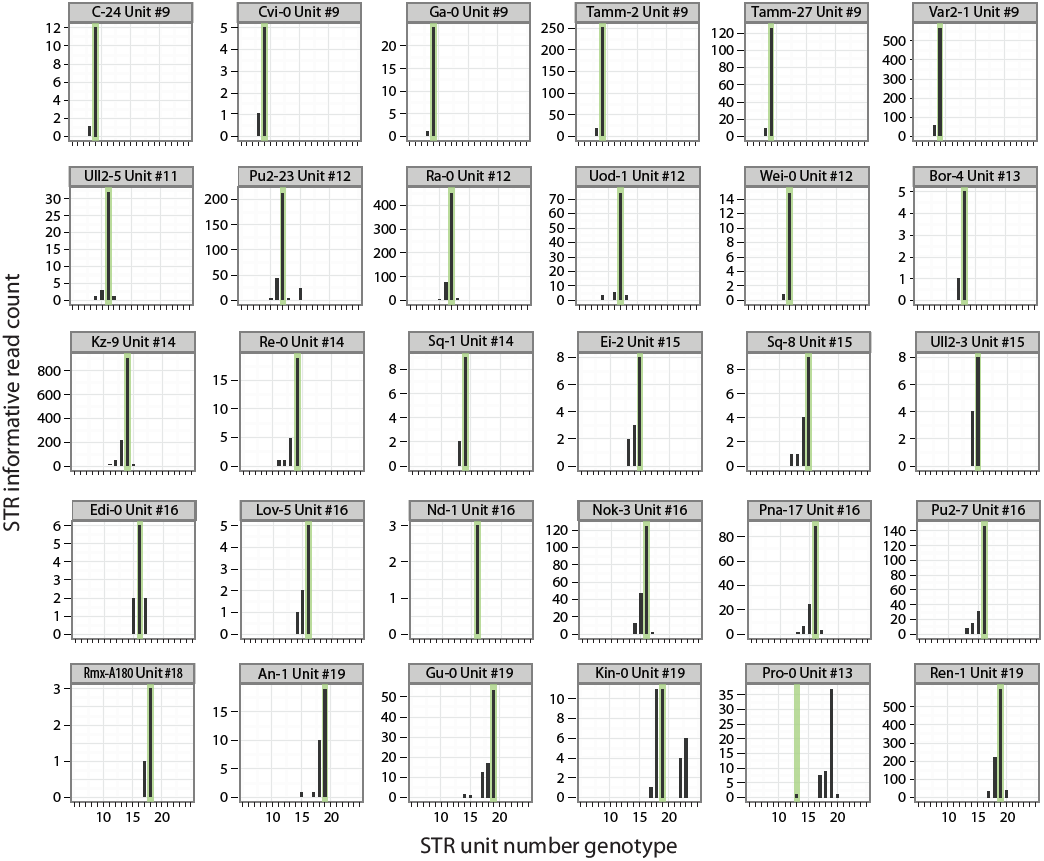
MIPSTR accurately determined germ-line *ELF3*-STR unit number on a population-scale across genetically diverse A. thaliana strains. Histograms of raw read counts across 30 accessions. STR unit number as determined by Sanger sequencing is indicated in green. Using tag-defined read groups, the Kin-0 *ELF3* STR genotype can be resolved to the known STR genotype even with comparatively few total reads. MIPSTR clearly calls STR unit number 19 for Pro-0. Note that different individuals of the same strain were analyzed with MIPSTR and Sanger-sequencing, which may explain the discrepancy.

The unit number, unit length, and purity of a given STR locus in a high-quality reference genome predict its variation across individuals (Legendre et al. 2007). STRs with high unit number, short unit length, and high purity are typically highly variable. With population-scale STR genotypes in hand, we addressed how well predicted variation of STRs (VARscore) (Legendre et al. 2007) correlated to observed variation across *A. thaliana* strains.

In general, VARscore correlated well with observed variation across STRs (r=0.68, Fig. 5), a substantially better agreement than previously observed (Duitama et al. 2014). However, this correlation was substantially weaker among coding STRs (r=0.46) than among non-coding STRs (r=0.75). This discrepancy suggests that sequence characteristics alone do not suffice to predict whether coding STRs vary on a population-scale. Coding STRs are more likely to be functionally important, and thus are less subject to the “neutral model” of the VARscore prediction.

**Figure 5.**
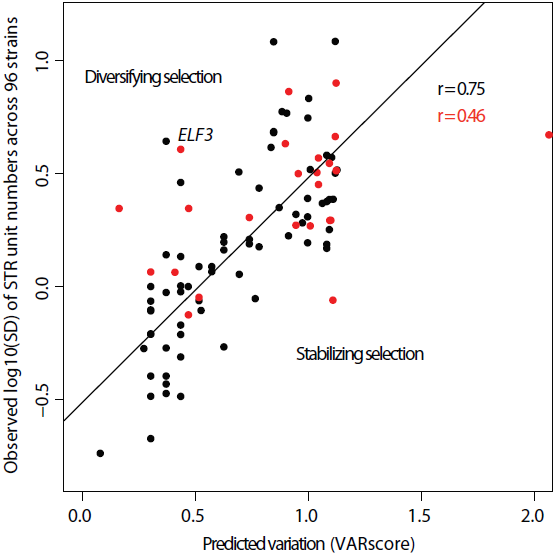
Observed and predicted STR variation showed greater correlation for non-coding STRs than coding STRs. The correlation between the observed log_10_ of the standard deviation of STR unit number across strains (y-axis) and the VARscore (x-axis), which predicts STR variation from sequence characteristics. Black points are non-coding STRs, red points are coding STRs. Outliers may indicate functional importance (*ELF3* STR is indicated).

Deviation of predicted STR variation (*i.e.* VARscore) from observed variation may thus hold information about STR function and selective pressures acting upon it. Specifically, STRs that are observed to be more variable than predicted may be under diversifying selection whereas those STRs that are observed to be less variable than predicted may be functionally constrained and under purifying selection (Press et al. 2014). For example, the STR in the gene *ELF3* is highly variable across strains, ranging from 7 units to as many as 29 units in a set of strains previously analyzed by Sanger sequencing (Undurraga et al. 2012). The phenotypes associated with variation in the *ELF3* STR change dramatically in different genetic backgrounds, suggesting co-evolution of the *ELF3-*STR with epistatically interacting loci (Undurraga et al. 2012). Given this STR’s strong background-dependent phenotypes, it is likely under diversifying selection and, correspondingly, it is much more variable than predicted (Fig. 5).

A complementary approach for identifying STRs with important function in modulating phenotype is genome-wide association of STR genotypes with phenotypes. The standard statistical methods for associating genotype with phenotype were developed for common, biallelic SNVs (Hayes 2013). STRs are typically multiallelic and often involved in epistatic interactions, both of which make it difficult to associate STR genotype with phenotype using standard methods (Press et al. 2014). Nevertheless, we performed a naïve association analysis to determine whether STR variation across strains was associated with well-characterized phenotypes (Atwell et al. 2010). We used the one-way analysis of variance (ANOVA) to detect associations between STR loci and phenotypes following previous studies (Mackay et al. 2012), modeling STR alleles as factors to avoid assumptions of linearity (Press et al. 2014). To minimize spurious associations, we dropped STRs that were typed in fewer than 10 strains from this analysis, and for each STR we dropped all strains carrying alleles present in fewer than three strains (rare alleles). We identified 124 significant associations involving 27 STRs and 41 phenotypes at a 1% false discovery rate (**Supplemental Table 3**). However, an important caveat is that this analysis did not consider population structure, which is another challenge given the different evolutionary trajectories of SNVs and STRs (Willems et al. 2014).

Our MIP-based approach can easily be scaled to thousands of targets; the human exome MIP set targets ∼55,000 loci (Turner et al. 2009). Over 2000 STR loci are accessible by MIPSTR in *A. thaliana,* and many more accessible STR loci exist in humans (Duitama et al. 2014; Guilmatre et al. 2013; Willems et al. 2014; Guilmatre et al. 2013; Molla et al. 2009). Our preliminary results, considering only a fraction of the accessible *A. thaliana* STR loci, highlight the promise of STRs to contribute to the variation and heritability of quantitative traits (Press et al. 2014).

## MIPSTR has potential to sensitively detect heterozygous and somatic STR unit number alleles

To determine the sensitivity with which MIPSTR detects heterozygous and somatic alleles, we mixed DNA of two divergent *A. thaliana* strains, Col-0 and Landsberg (Ler), in known ratios before MIPSTR capture and sequencing (Fig. 6). Of the 100 STR loci, 56 differed in STR unit number genotypes between Col-0 and Ler, and hence their relative presence across mixtures could be detected by MIPSTR. To assess the relative proportions of STR alleles within each mixture, we determined the number of tag-defined read groups for which the majority of reads supported either the Col-0-specific STR unit number or the Ler-specific STR unit number. This measure, however, is confounded by unequal coverage between libraries. More deeply sequenced libraries will represent a higher number of capture events per target and hence be more likely to identify rare STR alleles (*i.e*. somatic events). To account for variation in number of supporting tag-defined read groups per locus, we performed bootstrap resampling of the modes of the tag-defined read groups at each locus in each library 1000 times, while measuring the proportion of bootstrap samples in which the Col-0 allele was detected. Applying this method to our mixing experiment, the agreement between predicted and observed probabilities of observing Col-0 STR alleles was striking. For example, when we mimicked a “heterozygous” state with a 1:1 Col-0/Ler mixture we observed the Col-0 allele nearly 100% of the time. This agreement of predicted and observed probabilities held across all mixtures (Fig. 6), indicating that MIPSTR sensitively detects rare alleles. Mixing 1 part Col-DNA into 999 parts Ler-DNA, we were able to detect the Col-alleles at half of the 56 loci.

**Figure 6.**
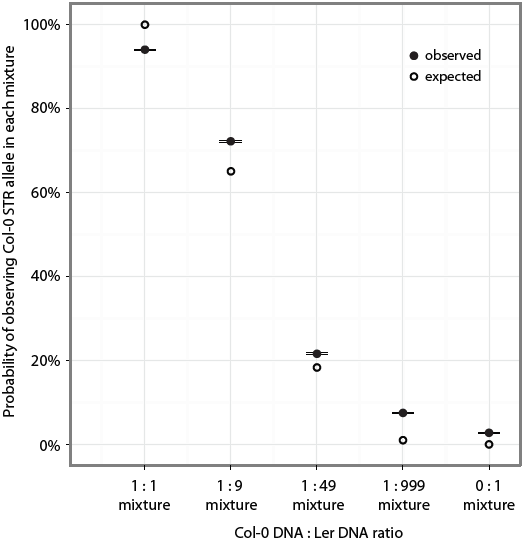
MIPSTR detects low frequency STR alleles. X-axis, tested mixtures of Ler and Col-0 DNA. Y-axis, probability of detecting Col-0 STR alleles. Closed circles are observed probability of observing Col-0 STR alleles (standard error is indicated, black lines); open circles are predicted probability of observing Col-0 STR alleles. To calculate the observed probability for each mixture, we re-sampled tag-defined read group modes supporting either the Col-0 or Ler allele at each STR locus 1000 times. The proportion of samples that carry the Col-0 allele was determined and averaged across all STR loci that differ between Ler and Col-0. To calculate the expected probability for each mixture, we assumed the known ratios of Col-0 and Ler STR alleles in each mixture and the probability of observing the Col-0 STR allele with ten observations.

STR instability at selected loci has been previously used as a measure of genome instability and is a hallmark of certain cancers (Kim et al. 2013; Boland et al. 1998). Our data suggest that MIPSTR has the potential to offer considerably greater resolution by assessing somatic STR variation genome-wide. To examine the potential of our method to detect decreased genome stability, we performed MIPSTR on *Atmsh2* mutant plants. This mutant carries an insertion in the *MSH2* gene, which is a crucial component of the DNA repair machinery. Indeed, a previous study, using a reporter system, found a ∼10% increase in dinucleotide STR somatic mutation events in this mutant (Golubov et al. 2010). We applied MIPSTR to three Col-0 plants and three *Atmsh2* plants. After eliminating STR loci with high technical error rates and loci without information for both strains, we compared the average number of STR alleles per locus with bootstrap resampling as described above. Instead of assessing two alleles, those of Col-0 and Ler as in the mixtures, we counted all alleles supported by at least one tag-defined read group in the resampling procedure. Compared to Col-0, the *Atmsh2* plants showed a 4.7% increase in average STR alleles across loci (p < 2.2E-16, Wilcoxon test, **Supplemental Fig. 4A**). Removing the two most overrepresented Col-0 and *Atmsh2* libraries, (*i.e.* with many more tag-defined read groups represented), resulted in an even larger difference between Col-0 and *Atmsh2*, with a 10.6% increase in *Atmsh2* mutants’ average STR alleles across all tested loci (p < 2.2E-16, Wilcoxon test, **Supplemental Fig. 4B**). This result is particularly remarkable considering that these loci were not optimized with respect to those most likely to exhibit somatic variation. Such optimization is readily possible with MIPSTR – for example, by applying MIPSTR to long non-coding dinucleotide STRs, which are far more prone to unit number mutation and hence somatic error. By combining such a specifically designed set of smMIPs (*i.e.* targets) for detecting somatic STR variation with deep sequencing, MIPSTR may be capable of identifying much more subtle increases in genome instability.

## Discussion

The potential of STR variation to contribute to phenotypic variation and heritability of complex traits is increasingly recognized (Press et al. 2014). To realize this potential, several recent efforts, relying on either analytical or experimental innovation, have made progress towards the ascertainment of accurate STR genotypes on a population-scale (Cao et al. 2014; Duitama et al. 2014; Guilmatre et al. 2013; Gymrek et al. 2012; Highnam et al. 2013). However, the STR-specific challenges for accurate genotyping – mappability and high amplification stutter – were only partially addressed. Here, we resolve these challenges by capturing STRs with single-molecule Molecular Inversion Probes that allow detection of many independent capture events of the same STR across many DNA molecules (Hiatt et al. 2013). Specifically, we resolve the mappability challenge by using targeted capture and locus-specific synthetic reference sequences. We resolve the challenge of inherently high technical error in STR amplification by examining many tag-derived read groups for each STR locus. STR unit number variation within a tag-defined read group results from amplification stutter. In contrast, STR unit number variation among tag-defined read groups has the potential to detect genomic duplications, heterozygosity, and somatic variation. We show that MIPSTR is capable of distinguishing these crucial sources of STR variation within samples.

Previous studies relied on amplification of haploid or homozygous genomes to estimate technical error for STR-containing sequencing reads (Guilmatre et al. 2013; Gymrek et al. 2012; Highnam et al. 2013); this approach is confounded by somatic variation and high STR mutation rates. MIPSTR offers an experimental avenue for empirically ascertaining technical error for many types of STRs. Notably, we observed dramatic differences in technical error even among the 100 trinucleotide and hexanucleotide STRs tested here. With larger numbers and more types of STRs, one may derive more precise predictions of sequencing error based on sequence composition, length, genomic position and other features.

However, even in this proof-of-principle study some patterns emerged that inform our understanding of the mutability of STRs. First, as others have seen, the most common technical error we observed was the loss of one STR unit (STR variation within tag-defined read groups) (Guilmatre et al. 2013). The loss of one STR unit was also the most common somatic event (STR variation observed among tag-defined read groups). As STR variation within a tag-defined read group exclusively derives from amplification stutter, we speculate that the somatic loss of one STR unit similarly derives from amplification errors during replication, rather than errors in DNA recombination or repair. Second, as anticipated by previous studies (2007 Legendre GR paper), longer STRs showed both increased technical and somatic error. Third, comparing predicted (based on neutral models) to observed variation in STR unit number, we found a stronger correlation for non-coding STRs than coding STRs, consistent with greater selective pressures on the latter, and suggesting that deviations from expected STR variation may hold information about an STR’s functional importance.

Although the immediate application of MIPSTR is in accurately assessing germ-line STR variation, we also emphasize our method’s potential to sensitively detect somatic STR variation. Somatic STR variation, better known as microsatellite instability (MSI), has a long history as a biomarker for certain colorectal cancers, more recently also for endometrial cancers (Boland et al. 1998; Kim et al. 2013). In fact, a recent study used exome sequencing data (∼20X coverage, 100 bp reads, compare with Figure S1) to assess MSI in colorectal and endometrial tumor and matched normal samples (Kim et al. 2013). Using only STR-spanning reads, this study called an MSI event at a given STR locus by comparing STR unit number distributions between tumor and matched normal samples, controlling for technical error with the STR variation observed in normal samples. As we show, comparing read distributions is vulnerable to differences in coverage and requires normalization by bootstrap resampling. MIPSTR eliminates the need to compare distributions of “normal” and ‘tumor’ samples to correct for technical error because MIPSTR calls both germ-line STR genotype and somatic STR variation in a given sample.

Although the STR loci that we targeted were not optimized for somatic events, MIPSTR detected the Col-0 STR alleles even in a 1:999 mixture of Col-0 and Ler-DNA. Moreover, using MIPSTR we observed a substantial increase of somatic events in a plant mutant deficient in DNA repair. MIPSTR can readily test and identify panels of 100-500 STR loci that are particularly unstable and prone to many somatic mutation events – for example by testing longer and less complex STRs such as di-or mononucleotides.

Beyond cancer genomics, at a population-scale, somatic variation and its occurrence across tissues, developmental stages, and in response to environmental perturbations has remained largely inaccessible due to the prohibitive costs of ultra-deep and single-cell sequencing (Baslan et al. 2012; Navin et al. 2011). As STRs are highly mutable, they are arguably the best biomarkers to detect even subtle perturbations of genome stability. We suggest that MIPSTR in combination with STR panels optimized for somatic variation has great promise to detect even subtle decreases in genome stability. We and others have previously proposed that subtly decreased genome stability may precede or coincide with many disease processes and may increase the penetrance of disease risk alleles (Queitsch et al. 2012; Heng 2010; Poduri et al. 2013). MIPSTR offers an approach to empirically test this hypothesis. Compared to single cell sequencing (Baslan et al. 2012; Navin et al. 2011). MIPSTR also offers a cost-and labor-efficient alternative for assessing the genetic heterogeneity of tumors, which is clinically relevant for disease treatment and prognosis (Fox et al. 2013; Schmitt et al. 2012)

Finally, we emphasize that MIPSTR is readily scalable: by simply targeting all STR loci in its size range, our method can provide genome-wide assessment of STR variation; by sequencing more deeply for an optimized panel of STR loci our method can provide information about somatic variation. MIPSTR is applicable to any organism with a high-quality reference genome, including humans. In the future, applying MIPSTR across populations of diverse species will contribute to fulfilling the long overdue promise of STR variation for explaining trait heritability.

## Methods

### smMIP capture reagent design

Each smMIP is an 80 bp oligonucleotide with a 40 bp common backbone flanked by an extension arm of 16-20 bp and a ligation arm of 20-24 bp. These unique arms specifically hybridize to flanking regions of STR loci for a gap-fill of 200 bp. Included in the 40 bp of the common backbone are 12 random nucleotides, the degenerate tag, generating ∼12^4^ = 1.67 × 10^6^ unique sequences per MIP. The MIPs were designed for 102 STRs across the *A. thaliana* genome (**Supplemental Table 1**).

These MIPs were procured individually by column-synthesis on an 100 nmol scale with standard desalting purification (at a cost of ∼$32 per MIP). Once purchased, one has effectively an infinite MIP supply allowing for millions of capture reactions, justifying the considerable upfront MIP cost. Cost per MIP is significantly lower when ordering less MIP without purification (25 nmol/$7.20 per MIP) (Hiatt et al. 2013).

MIPs were pooled at equal molarity and mixed with the target at 200-fold molar excess. The results of the first capture reaction in the Col-0 reference genome, specifically the distribution of read counts from each MIP, were used to adjust MIP concentrations. We increased the concentration of the lowest performing MIPs (28, fewest number of reads) 100-fold; concentration of the next lowest performing group of MIPS (43) was increased 10-fold.

## Capture and Library Construction

DNA was extracted from rosette leaves of individual 20-day-old *A. thaliana* plants using DNeasy Plant Maxi Kit (Qiagen). DNA was cleaned up and concentrated with Amicon Ultra Centrifugal Filter Units (Millipore).

Capture procedures were modified from previous protocols (O’Roak et al. 2012; Hiatt et al. 2013). 750 ng genomic DNA was mixed with 2 pmol smMIP mixture (starting concentration before adjustment for low performing MIPs), 1.5 µl 10X Ampligase buffer, and molecular biology grade water to a total volume of 15 µl. For hybridization, these mixtures were incubated in a thermocycler with a heated lid for 10 minutes at 95°C followed by 48 hours at 55°C. After hybridization, we added 2.5 pmol dNTPs (TaKaRa), 1 unit Ex Taq polymerase (TaKaRa), 0.5 µl 10x Ampligase buffer, 60 units Ampligase DNA ligase (Epicentre) and molecular grade water to an added volume of 5 µl per mixture. The extension phase was carried out at 60°C for an hour. After gap-fill and ligation, the mixtures were cooled to 37°C for two minutes. We then added 40 units of Exonuclease I (NEB) and 200 units of Exonuclease III (NEB) for a total reaction volume of 19 µl. To digest uncircularized and excess genomic DNA, we incubated these mixtures at 37°C for 15 minutes, and then denatured the enzymes at 92°C for two minutes.

## Library construction, purification, and pooling

To create sequencing libraries, we amplified the capture reactions using a common forward primer and an indexed reverse primer. We mixed 5 µl capture reaction with 12.5 pmol dNTPs (TaKaRa), 5 ul 10X Ex Taq buffer, 25 micromoles forward primer, 25 micromoles reverse primer, 1 unit Ex Taq polymerase (TaKaRa), and molecular biology grade water to a total reaction volume of 50 µl. We performed an initial denaturation at 98°C for 10 seconds, followed by 28 cycles of 10 seconds at 98°C, 30 seconds at 58°C and 12 seconds at 72°C. The final extension was for 3 minutes at 72°C. PCR products were pooled as equal volumes per sample or according to gel image quantification to get approximately equal representation. We then cleaned up the pooled PCR products using Ampure XP beads (Agencourt) at 1.8X according to manufacturer’s recommendations.

## Sequencing and primary analysis

Samples were sequenced using the Illumina MiSeq v2 platform according to the manufacturer’s instructions with custom sequencing primers (Hiatt et al. 2013). To improve cluster generation for these low complexity STR libraries, we spiked in Phi-X or whole genomic DNA libraries at 10-20%. We collected one 250 bp forward read to determine sequence of the ligation arm and STR target locus, one 50 bp reverse read to determine the sequence of the degenerate tag and extension arm, and one 8 bp read to determine the sample index sequence.

The MiSeq software sorted by index read to separate pooled libraries.

## Mapping and STR genotype calling

For each target STR locus, we created a synthetic reference of 100 “chromosomes,” which consisted of the Col-0 reference target sequence with 1 to 100 pure STR units (no SNVs). We sorted reads by the first 16 bp of the ligation targeting arm allowing three mismatches and then used the bwasw alignment mode of the *bwa* aligner (Li and Durbin 2009) to map the reads to the locus-specific synthetic reference. For a given read, if the A-score of its alignment to a specific synthetic “chromosome” was ≥180, we called the STR unit number of this “chromosome” for this read. Below this A-score, the read was discarded. When the sequence read ended within the STR (presumably due to a large expansion of STR units) but still mapped with an acceptable A-score, we called the genotype as ≥ the unit number of the “chromosome” to which the read aligned. In this way, MIPSTR can yield information about STR unit number expansions in a given individual even in the absence of STR-spanning reads. Here, these “≥” calls were not used in further analyses such as association or calculation of variation.

We then sorted the STR genotype calls by the degenerate tag on the paired reverse read from which they derived. We required an exact match of the 12 bp degenerate tag for reads to be grouped into a tag-defined read group. We then called the mode STR unit number of each tag-defined read group as the genotype of that DNA molecule. If we observed that more than one tag-defined read group supported an alternate STR allele, we considered it evidence of somatic variation.

## STR association with phenotypes

We used previously published data for 107 phenotypes collected for 96 *A. thaliana* strains (Atwell et al. 2010). We then proceeded to detect associations between each of these phenotypes and each variable STR locus within genotyped strains. For each test, we omitted strains from the analysis that were not phenotyped for the relevant trait or genotyped at the STR in question. We additionally removed from the analysis strains that carried STR alleles that were found in fewer than three strains total, to avoid confounding from rare alleles. We then performed one-way ANOVA to test the null hypothesis of no association between each STR and each phenotype, while treating each STR allele categorically. We chose to treat STR alleles categorically because assumptions of linearity in STR-phenotype associations are poorly founded in some cases (Undurraga et al. 2012; Press et al. 2014). Associations were accepted at a 1% false discovery rate (p = 1.48 * 10^−4^).

## Calculating technical error rates

To calculate the technical error rate of amplifying STR loci, we considered all tag-defined read groups for which a single STR unit number mode was supported by at least two reads. For these tag-defined read groups, we took the fraction of reads supporting unit numbers other than the mode and divided by the total number of reads. We averaged across all tag-defined read groups at a given locus for a technical error score between 0 and 1 representing the fraction of reads at a locus known to be error (**Supplemental Table 1**).

## Somatic allele counts

To compare the number of somatic events occurring in different individuals, we only considered STR loci with low technical error scores (below 0.2, **Supplemental Table 1**) and with information for all plants in the comparison. We used bootstrap resampling to account for sometimes vastly different read counts. For example, in the Col-0 and Ler mixing experiment, some mixture libraries had as few as ten tag-defined read groups at a given locus. Thus, we resampled ten modes from tag-defined read groups in these samples, counting the proportion of those samples in which the Col-0 unit number allele was present. In Col-0 versus *atmsh2* experiment, depth of coverage was much higher and hence we resampled 1000 modes of tag-defined read groups for each locus. For each sample, we calculated how many different STR unit number alleles were present and averaged across loci.

## Acknowledgements

This work was supported by grants from the National Human Genome Research Institute Interdisciplinary Training in Genomic Sciences (2T32HG35-16 to MOP, T32 HG00035 to KDC) and the National Institute of Health New Innovator Award (DP2OD008371 to CQ). We would like to thank Matt Rich, Josh Cuperus, Matthew Snyder, Akash Kumar, Joe Hiatt, Choli Lee, Rachel Youngblood, Giang Ong, Jacob Kitzman, and Queitsch lab members for helpful discussions.

## References

Atwell S, Huang YS, Vilhjálmsson BJ, Willems G, Horton M, Li Y, Meng D, Platt A, Tarone AM, Hu TT, et al. 2010. Genome-wide association study of 107 phenotypes in Arabidopsis thaliana inbred lines. Nature 465: 627–631.

Baslan T, Kendall J, Rodgers L, Cox H, Riggs M, Stepansky A, Troge J, Ravi K, Esposito D, Lakshmi B, et al. 2012. Genome-wide copy number analysis of single cells. Nat Protoc 7: 1024–1041.

Boland CR, Thibodeau SN, Hamilton SR, Sidransky D, Eshleman JR, Burt RW, Meltzer SJ, Rodriguez-Bigas MA, Fodde R, Ranzani GN, et al. 1998. A National Cancer Institute Workshop on Microsatellite Instability for cancer detection and familial predisposition: development of international criteria for the determination of microsatellite instability in colorectal cancer. Cancer Res 58: 5248–5257.

Bolton KA, Ross JP, Grice DM, Bowden NA, Holliday EG, Avery-Kiejda KA, Scott RJ. 2013. STaRRRT: a table of short tandem repeats in regulatory regions of the human genome. BMC Genomics 14:795.

Butler AP, Trono D, Coletta LD, Beard R, Fraijo R, Kazianis S, Nairn RS. 2007. Regulation of CDKN2A/B and Retinoblastoma genes in Xiphophorus melanoma. Comp Biochem Physiol Toxicol Pharmacol CBP 145: 145–155.

Cao J, Schneeberger K, Ossowski S, Günther T, Bender S, Fitz J, Koenig D, Lanz C, Stegle O, Lippert C, et al. 2011. Whole-genome sequencing of multiple Arabidopsis thaliana populations. Nat Genet 43: 956–963.

Cao MD, Tasker E, Willadsen K, Imelfort M, Vishwanathan S, Sureshkumar S, Balasubramanian S, Bodén M. 2014. Inferring short tandem repeat variation from paired-end short reads. Nucleic Acids Res 42: e16.

Chen K, Wallis JW, McLellan MD, Larson DE, Kalicki JM, Pohl CS, McGrath SD, Wendl MC, Zhang Q, Locke DP, et al. 2009. BreakDancer: an algorithm for high-resolution mapping of genomic structural variation. Nat Methods 6: 677–681.

Duitama J, Zablotskaya A, Gemayel R, Jansen A, Belet S, Vermeesch JR, Verstrepen KJ, Froyen G. 2014. Large-scale analysis of tandem repeat variability in the human genome. Nucleic Acids Res 42: 5728–5741.

Eckert KA, Hile SE. 2009. Every microsatellite is different: Intrinsic DNA features dictate mutagenesis of common microsatellites present in the human genome. Mol Carcinog 48: 379–388.

Fondon JW 3rd, Garner HR. 2004. Molecular origins of rapid and continuous morphological evolution. Proc Natl Acad Sci U S A 101: 18058–18063.

Fox EJ, Prindle MJ, Loeb LA. 2013. Do mutator mutations fuel tumorigenesis? Cancer Metastasis Rev 32: 353–361.

Gan X, Stegle O, Behr J, Steffen JG, Drewe P, Hildebrand KL, Lyngsoe R, Schultheiss SJ, Osborne EJ, Sreedharan VT, et al. 2011. Multiple reference genomes and transcriptomes for Arabidopsis thaliana. Nature 477: 419–423.

Gatchel JR, Zoghbi HY. 2005. Diseases of unstable repeat expansion: mechanisms and common principles. Nat Rev Genet 6: 743–755.

Gemayel R, Cho J, Boeynaems S, Verstrepen KJ. 2012. Beyond junk-variable tandem repeats as facilitators of rapid evolution of regulatory and coding sequences. Genes 3: 461–480.

Golubov A, Yao Y, Maheshwari P, Bilichak A, Boyko A, Belzile F, Kovalchuk I. 2010. Microsatellite instability in Arabidopsis increases with plant development. Plant Physiol 154: 1415–1427.

Grimm D, Hagmann J, Koenig D, Weigel D, Borgwardt K. 2013. Accurate indel prediction using paired-end short reads. BMC Genomics 14: 132.

Guilmatre A, Highnam G, Borel C, Mittelman D, Sharp AJ. 2013. Rapid multiplexed genotyping of simple tandem repeats using capture and high-throughput sequencing. Hum Mutat 34: 1304–1311.

Gymrek M, Golan D, Rosset S, Erlich Y. 2012. lobSTR: A short tandem repeat profiler for personal genomes. Genome Res 22: 1154–1162.

Hajirasouliha I, Hormozdiari F, Alkan C, Kidd JM, Birol I, Eichler EE, Sahinalp SC. 2010. Detection and characterization of novel sequence insertions using paired-end next-generation sequencing. Bioinforma Oxf Engl 26: 1277–1283.

Hammock EAD, Young LJ. 2005. Microsatellite instability generates diversity in brain and sociobehavioral traits. Science 308: 1630–1634.

Hayes B. 2013. Overview of Statistical Methods for Genome-Wide Association Studies (GWAS). Methods Mol Biol Clifton NJ 1019: 149–169.

Heng HHQ. 2010. Missing heritability and stochastic genome alterations. Nat Rev Genet 11: 813.

Hiatt JB, Pritchard CC, Salipante SJ, O’Roak BJ, Shendure J. 2013. Single molecule molecular inversion probes for targeted, high-accuracy detection of low-frequency variation. Genome Res 23: 843–854.

Highnam G, Franck C, Martin A, Stephens C, Puthige A, Mittelman D. 2013. Accurate human microsatellite genotypes from high-throughput resequencing data using informed error profiles. Nucleic Acids Res 41: e32.

Kim T-M, Laird PW, Park PJ. 2013. The landscape of microsatellite instability in colorectal and endometrial cancer genomes. Cell 155: 858–868.

Legendre M, Pochet N, Pak T, Verstrepen KJ. 2007. Sequence-based estimation of minisatellite and microsatellite repeat variability. Genome Res 17: 1787–1796.

Li H, Durbin R. 2009. Fast and accurate short read alignment with Burrows-Wheeler transform. Bioinforma Oxf Engl 25: 1754–1760.

Mackay TFC, Richards S, Stone EA, Barbadilla A, Ayroles JF, Zhu D, Casillas S, Han Y, Magwire MM, Cridland JM, et al. 2012. The Drosophila melanogaster Genetic Reference Panel. Nature 482: 173–178.

Michael TP, Park S, Kim T-S, Booth J, Byer A, Sun Q, Chory J, Lee K. 2007. Simple sequence repeats provide a substrate for phenotypic variation in the Neurospora crassa circadian clock. PloS One 2: e795.

Molla M, Delcher A, Sunyaev S, Cantor C, Kasif S. 2009. Triplet repeat length bias and variation in the human transcriptome. Proc Natl Acad Sci U S A 106: 17095–17100.

Mularoni L, Guigó R, Albà MM. 2006. Mutation patterns of amino acid tandem repeats in the human proteome. Genome Biol 7: R33.

Navin N, Kendall J, Troge J, Andrews P, Rodgers L, McIndoo J, Cook K, Stepansky A, Levy D, Esposito D, et al. 2011. Tumour evolution inferred by single-cell sequencing. Nature 472: 90–94.

Nithianantharajah J, Hannan AJ. 2007. Dynamic mutations as digital genetic modulators of brain development, function and dysfunction. BioEssays News Rev Mol Cell Dev Biol 29: 525–535.

O’Dushlaine CT, Edwards RJ, Park SD, Shields DC. 2005. Tandem repeat copy-number variation in protein-coding regions of human genes. Genome Biol 6: R69.

O’Roak BJ, Vives L, Fu W, Egertson JD, Stanaway IB, Phelps IG, Carvill G, Kumar A, Lee C, Ankenman K, et al. 2012. Multiplex targeted sequencing identifies recurrently mutated genes in autism spectrum disorders. Science 338: 1619–1622.

Peixoto AA, Hennessy JM, Townson I, Hasan G, Rosbash M, Costa R, Kyriacou CP. 1998. Molecular coevolution within a Drosophila clock gene. Proc Natl Acad Sci U S A 95: 4475–4480.

Poduri A, Evrony GD, Cai X, Walsh CA. 2013. Somatic mutation, genomic variation, and neurological disease. Science 341: 1237758.

Press M, Carlson KD, Queitsch C. 2014. The overdue promise of short tandem repeat variation for heritability. http://biorxiv.org/lookup/doi/10.1101/006387 (Accessed July 10, 2014).

Qi J, Zhao F. 2011. inGAP-sv: a novel scheme to identify and visualize structural variation from paired end mapping data. Nucleic Acids Res 39: W567–575.

Queitsch C, Carlson KD, Girirajan S. 2012. Lessons from model organisms: phenotypic robustness and missing heritability in complex disease. PLoS Genet 8: e1003041.

Rosas U, Mei Y, Xie Q, Banta JA, Zhou RW, Seufferheld G, Gerard S, Chou L, Bhambhra N, Parks JD, et al. 2014. Variation in Arabidopsis flowering time associated with cis-regulatory variation in CONSTANS. Nat Commun 5: 3651.

Sawyer LA, Hennessy JM, Peixoto AA, Rosato E, Parkinson H, Costa R, Kyriacou CP. 1997. Natural variation in a Drosophila clock gene and temperature compensation. Science 278: 2117–2120.

Scarpino SV, Hunt PJ, Garcia-De-Leon FJ, Juenger TE, Schartl M, Kirkpatrick M. 2013. Evolution of a genetic incompatibility in the genus Xiphophorus. Mol Biol Evol 30: 2302–2310.

Schmitt MW, Prindle MJ, Loeb LA. 2012. Implications of genetic heterogeneity in cancer. Ann N Y Acad Sci 1267: 110–116.

Tae H, McMahon KW, Settlage RE, Bavarva JH, Garner HR. 2013. ReviSTER: an automated pipeline to revise misaligned reads to simple tandem repeats. Bioinforma Oxf Engl 29: 1734–1741.

Turner EH, Lee C, Ng SB, Nickerson DA, Shendure J. 2009. Massively parallel exon capture and library-free resequencing across 16 genomes. Nat Methods 6: 315–316.

Undurraga SF, Press MO, Legendre M, Bujdoso N, Bale J, Wang H, Davis SJ, Verstrepen KJ, Queitsch C. 2012. Background-dependent effects of polyglutamine variation in the Arabidopsis thaliana gene ELF3. Proc Natl Acad Sci U S A 109: 19363–19367.

Verstrepen KJ, Jansen A, Lewitter F, Fink GR. 2005. Intragenic tandem repeats generate functional variability. Nat Genet 37: 986–990.

Willems TF, Gymrek M, Highnam G, The 1000 Genomes Project, Mittelman D, Erlich Y. 2014. The Landscape of Human STR Variation. http://biorxiv.org/lookup/doi/10.1101/004671 (Accessed July 15, 2014).

Yu F, Sabeti PC, Hardenbol P, Fu Q, Fry B, Lu X, Ghose S, Vega R, Perez A, Pasternak S, et al. 2005. Positive selection of a pre-expansion CAG repeat of the human SCA2 gene. PLoS Genet 1: e41.

